# Computationally Guided Design of Novel Angiotensin Converting Enzyme Inhibitors

**DOI:** 10.1101/2022.04.27.489637

**Authors:** Jennifer D. Lam, Mary Riley, Justin B. Siegel

**Author notes:** Department of Chemistry, University of California, Davis, Davis, California, United States of America, Department of Biochemistry and Molecular Medicine, University of California, Davis, Davis, California, United States of America, Genome Center, University of California, Davis, Davis, California, United States of America.

## Abstract

Angiotensin-converting enzyme inhibitors (ACEI) such as Moexipril, Trandolapril, Ramipril, and Perindopril are used to treat hypertension. However, these drugs have undesired side effects. Using computational studies, two new novel drug candidates were designed with improved molecular interactions, ADMET properties, and docking scores compared to Moexipril. Homology analysis was done to determine the best animal model for preclinical studies, and it revealed that *Pan troglodytes* (chimpanzees) and *Mus musculus* (house mouse) were the best candidates.

## INTRODUCTION

Hypertension is a serious condition where blood pressure levels are constantly elevated.^1^ This can increase the risk of heart diseases, strokes, and heart attacks if it is not managed correctly.^1,2^ In 2018, 494,873 people died from hypertension in the United States.^1^ To combat this, pharmaceutical companies developed medications that inhibit an angiotensin-converting enzyme (ACE).^3^

Angiotensin-converting enzyme (ACE) plays a key role in the renin-angiotensin-aldosterone system that regulates blood pressure, which makes the enzyme a key target for inhibition.^3^ ACE transforms angiotensin I to angiotensin II which causes blood vessels to become constricted, making it more difficult for the heart to pump blood.^4,5^ The change of angiotensin I to II happens when histidine and leucine on the carboxyl-terminus are cleaved off of angiotensin I. The vasodilator mediator, bradykinin, loses its ability to dilate blood vessels because it becomes inactivated.^4^ The current solution to this problem is a class of drugs called angiotensin-converting enzyme inhibitors (ACEIs).^5^

There are many known ACEIs such as Trandolapril, Ramipril, Moexipril, and Perindopril that are successful in lowering blood pressure.^5^ The main role of blood pressure medications is to block the conversion of angiotensin I to angiotensin II by inhibiting ACE.^5^ ACEIs allow blood vessels to dilate and helps the body maintain its osmolarity by removing water.^6,7^ Maintaining normal blood pressure levels with ACEIs has contributed to the reduced risk of heart attacks, strokes, and heart diseases.^6^ Even though these drugs have been approved for many decades, there are still many concerns relating to their pharmacodynamic and pharmacokinetic properties. For example, some hypertension medication can be metabolized into reactive toxicophores by cytochrome P450 enzymes.^8,9^ Drugs such as Trandolapril and Ramipril can have negative effects on elderlies with compromised renal systems.^10,11^ ACEIs taken with non-steroidal anti-inflammatory drugs (NSAIDs) can cause acute renal failure.^10,11^ Trandolapril and Ramipril can also increase the chance of fetal morbidity and death in the second and third trimester of pregnancy.^10,11^ Some of the common side effects of taking ACEIs are dizziness, fatigue, headaches, cough, hypotension, and stomach problems.^6^

With the purpose of designing better ACEIs, computational studies were conducted to design two new drug candidates starting with the identification of the pharmacophore. Researchers have identified the pharmacophore of ACEIs as succinyl-L-proline (Figure 1).^13^ It is the pharmacophore because all ACE inhibitor drugs contains a proline, a carbonyl, an alkyl chain, and a carboxyl group.^10,11,13,14^ Therefore, the pharmacophore was used as a model for designing new drug candidates. The first novel drug candidate was designed using a computational program while the second was designed using chemical intuition. It was found that both candidates were potential leads for new ACEIs because they had an improvement in their docking scores, bonding interactions, and ADMET (absorption, distribution, metabolism, excretion, and toxicity) properties as compared to Moexipril. A homolog analysis was also used to predict which organisms would be a good model for preclinical studies.

**Figure 1.**
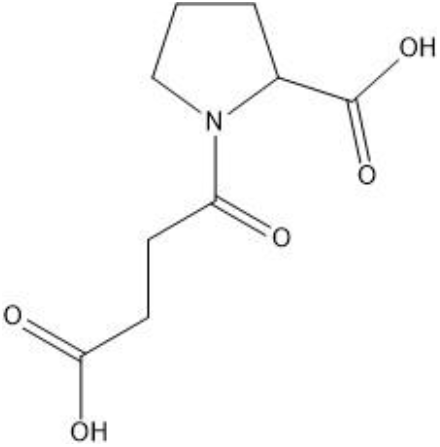
Structure of succinyl-L-proline, pharmacophore of ACE inhibitor drugs.

## METHODS

The crystal structure of human angiotensin-converting enzyme in complex with lisinopril (PDB ID: 1O86) was obtained from the RCSB Protein Data Bank website.^15^ Pymol was used to visualize the interactions between the ligand and amino acids of the enzyme.^16^

Make Receptor was used to define the binding site from the PDB code that was obtained.^17^ GaussView was used to construct molecules and Gaussian09 was used to optimize them.^18,19^ VIDA was also used to visualize and analyze the drug molecules, as well as the active site.^20^ vBrood was used to generate bioisosteric replacements to find novel drug molecules with better binding affinity and ADMET properties.^21^

The OpenEye FILTER was used to screen compounds based on their functional groups and physical properties.^22^ This determined the absorption, distribution, metabolism, elimination, and toxicity (ADMET) properties.^22^ OpenEye OMEGA was used to generate conformer libraries of the drug molecules.^23^ Conformer libraries were docked into the active site using FRED, which allows OpenEye to generate a docking score.^24^ BLAST was used to search for homologous proteins.^25^ Lastly, Jalview was used for sequence alignment of homologous proteins.^26^

## RESULTS AND DISCUSSION

### Evaluations of known ACEI drugs

To figure out which drug should be improved, four known ACEIs were examined (Trandolapril, Ramipril, Moexipril, and Perindopril). Docking scores, ADMET properties, and active site interactions were evaluated (Table 1, Figure 2). Ramipril had the highest docking score at −10.64 followed by Trandolapril at −10.30, and Perindopril at −8.86. Moexipril had the worst docking score at −5.59, which made it a good candidate for improvement because it had the lowest binding affinity to the active site. Additionally, it has other properties that make it a better ACEI than the other three drugs.

**Table 1.** ADMET properties and docking scores of ACEIs.

| Name | Molecular Weight (g/mol) | Lipinski H-bond Donor | Lipinski H-bond Acceptor | XLogP | Number of Chiral Centers | Docking Score |
| --- | --- | --- | --- | --- | --- | --- |
| Moexipril | 498.57 | 1 | 9 | 0.87 | 3 | -5.59 |
| Trandolapril | 430.54 | 1 | 7 | 1.37 | 5 | -10.30 |
| Ramipril | 416.51 | 1 | 7 | 0.88 | 5 | -10.64 |
| Perindopril | 368.47 | 1 | 7 | 0.49 | 5 | -8.86 |

**Figure 2.**
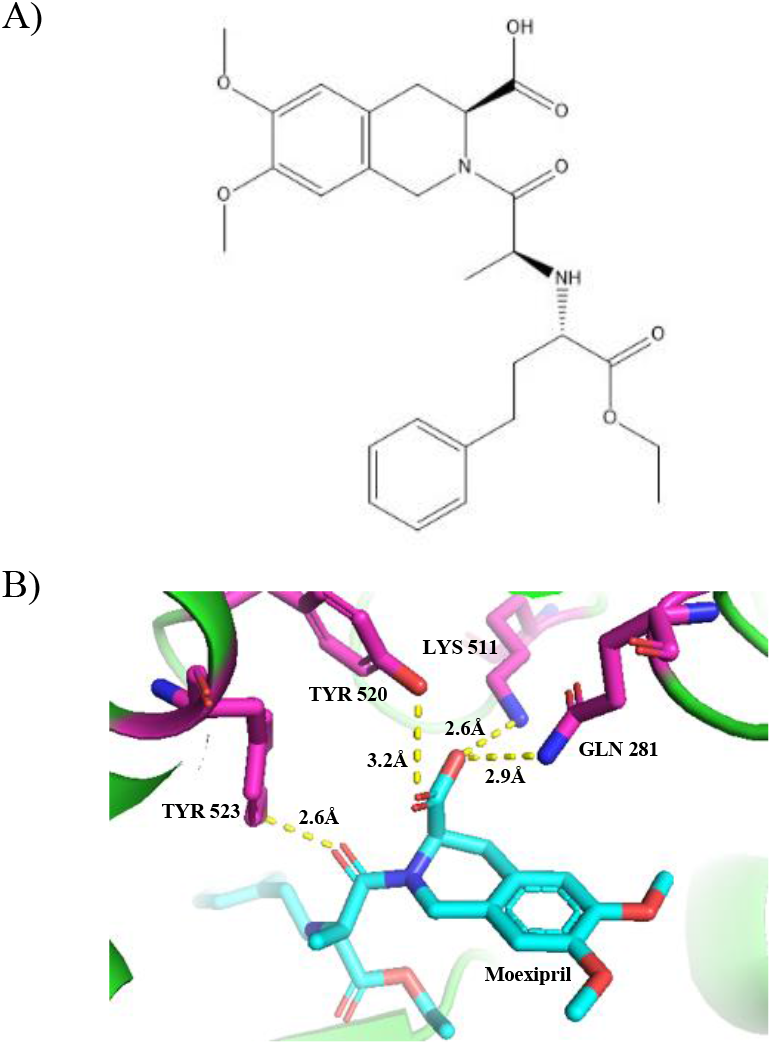
A) 2D structure of Moexipril. B) Moexipril (cyan) interactions with GLN 281, LYS 511, TYR 520, and TYR 523 residues (pink).

The ADMET screening showed no known toxicophores, aggregators, or violations of Lipinski Rule of 5 for all four drug molecules. However, the molecular weight of Moexipril came close to breaking Lipinski Rule at 498.57 g/mol. Comparing Trandolapril, Ramipril, and Perindopril, they all had one Lipinski hydrogen bond donor and seven Lipinski hydrogen bond acceptors. Similarly, Moexipril had one hydrogen bond donor, but also two additional hydrogen bond acceptors. This made it a better candidate with the possibility to make more hydrogen bonding and/or electrostatic interactions with residues in the active site. Additionally, Moexipril had three chiral centers while the others had five. It’s better to have fewer chiral centers to reduce the amount of possible off target effects.

Pymol showed that Moexipril had strong hydrogen bonds with residues GLN 281, LYS 511, TYR 520, and TYR 523 at distances of 2.9Å, 2.6Å, 3.2Å, and 2.6Å respectively (Figure 2B). Based on the comparison between the ACEIs, Moexipril was the most promising drug for additional improvements because it had the worst docking score, even though it had better ADMET properties. Therefore, two novel drug candidates were made using bioisosteric replacements to create additional molecular interactions and improve on the docking score and ADMET properties of Moexipril.

### Computationally Driven Design of Candidate 1

Candidate 1 (Figure 3A) was generated by loading Moexipril into vBrood. The benzene ring and ether groups attached to it did not have any interactions with the active site, so they were modified to improve binding affinity. The generated molecule had the best docking score when the ether groups were replaced with hydroxyl groups and one of the carbons on the benzene ring was replaced with a nitrogen. Additionally, the nitrogen on the adjacent cyclohexane was removed and double bonds were added. All these modifications created additional hydrogen bond donors and acceptors.

**Figure 3.**
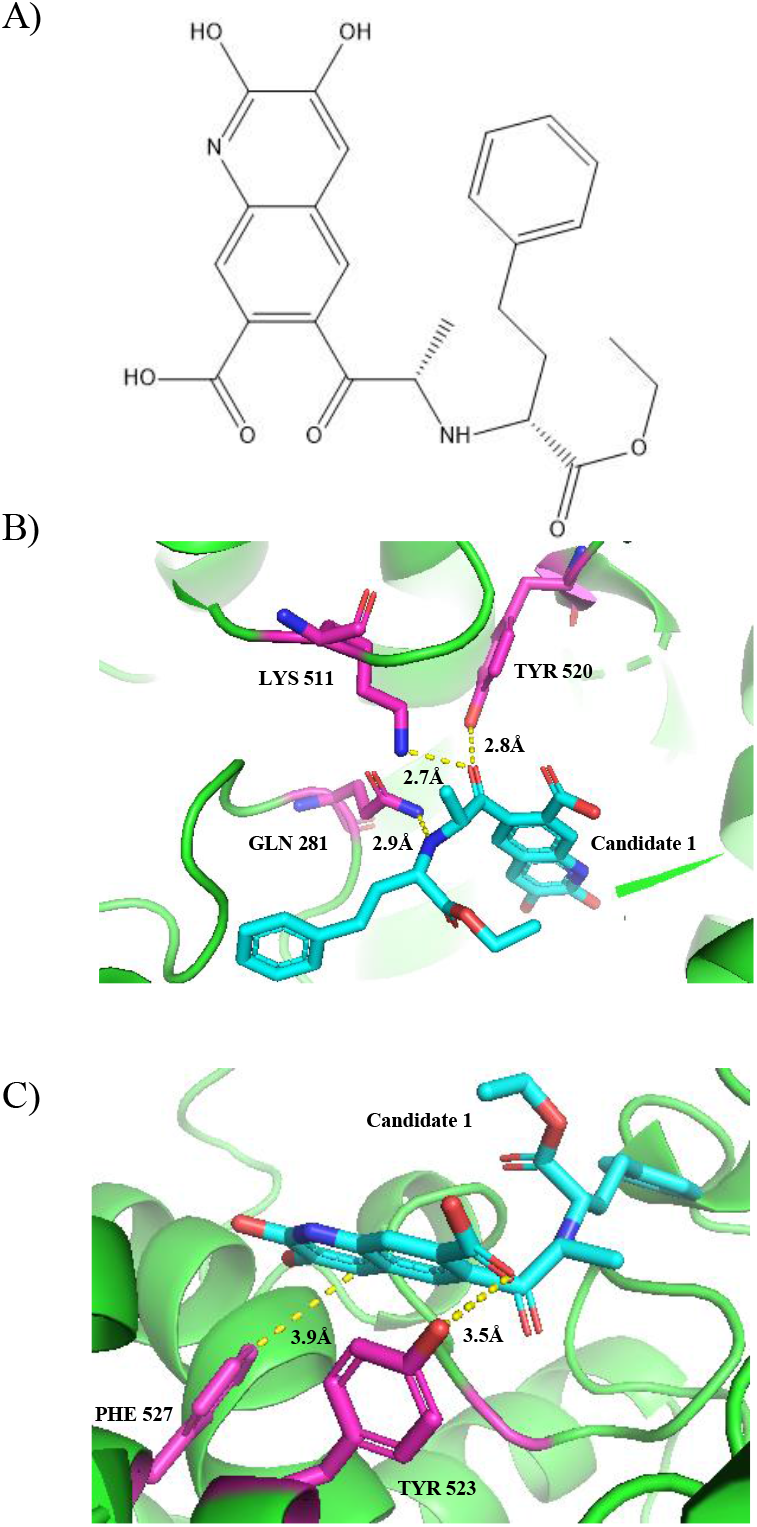
A) 2D structure of Candidate 1. B) Candidate 1 (cyan) interactions with GLN 281, LYS 511, and TYR 520 residues (pink). C) Candidate 1 (cyan) interactions with TYR 523, and PHE 527 residues (pink).

Candidate 1 had a docking score of −9.60 (Table 2), which was a 71.7% improvement compared to the original Moexipril molecule. Candidate 1 docked differently into the active site because of different molecular interactions between the residues. The modified hydroxyl functional groups did not have any new interactions with the residues. However, it shifted the molecule in the active site, creating new interactions similar to T-shape pi-pi stacking. All the interactions are shown in Figures 3B and 3C. TYR 520 made a 2.8Å hydrogen bond with a different carbonyl group, which shifted the molecule deeper into the active site. A new 2.7Å hydrogen bond interaction was also made between the carbonyl group of Candidate 1 and LYS 511. GLN 281 and TYR 523 both made new hydrogen bonds with Candidate 1 at 2.9Å and 3.5Å respectively. Even though some hydrogen bonds became longer and weaker, others became shorter and stronger. An additional new interaction made a 3.9Å T-shape pi-pi stacking interaction between PHE 527 and Candidate 1. These interactions contributed to the more negative docking score and changed how the molecule was positioned inside of the active site.

**Table 2.** ADMET properties and docking score of Candidate 1 and Candidate 2.

| Name | Molecular Weight (g/mol) | Lipinski H-bond Donor | Lipinski H-bond Acceptor | XLogP | Number of Chiral Centers | Docking Score |
| --- | --- | --- | --- | --- | --- | --- |
| Candidate 1 | 466.48 | 3 | 9 | 2.86 | 2 | -9.60 |
| Candidate 2 | 440.45 | 5 | 9 | 1.34 | 2 | -10.78 |

Similar to the ADMET properties of Moexipril, Candidate 1 had no known violations of Lipinski Rule of 5, toxicophores, or aggregators. The molecular weight of the drug decreased to 466.48 g/mol, increasing its chance of getting absorbed. However, the XLogP increased to 2.86. There were two additional Lipinski hydrogen bond donors which increased the number of possible hydrogen bonds or electrostatic interactions the drug can have with the active site. The number of chiral centers also decreased to two, decreasing the probability of having adverse effects. Based on the docking score, molecular interactions, and ADMET properties, Candidate 1 is expected to be a promising novel drug candidate.

### Chemical Intuition Driven Design of Candidate 2

Candidate 2 (Figure 4) was designed based off Candidate 1 by replacing the two oxygen atoms on the ethyl ester group with two hydroxyl groups. The hydroxyl groups can act as a hydrogen bond acceptor or donor. Candidate 2 had a docking score of −10.78 (Table 2), which is a 92.8% improvement compared to the original drug, Moexipril.

**Figure 4.**
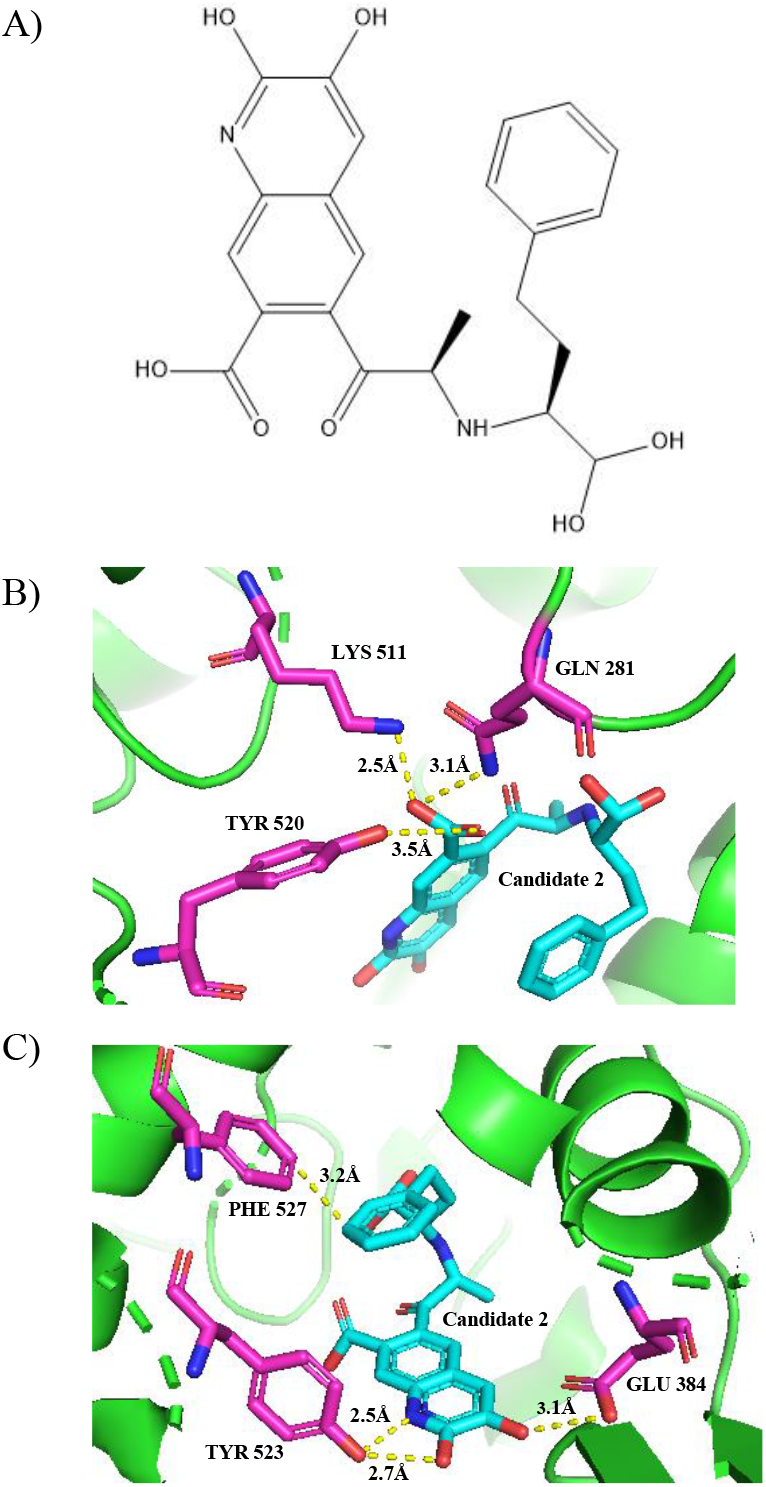
A) 2D structure of Candidate 2. B) Candidate 2 (cyan) interactions with GLN 281, LYS 511, and TYR 520 residues (pink). C) Candidate 2 (cyan) interactions TYR 523, GLU 384, and PHE 527 residues (pink).

Most of the molecular interaction distances remain similar but some changed when comparing to the original drug. These interactions with the active site are shown in Figure 4B and 4C. The hydrogen bonds made with GLN 281 and TYR 520 became slightly longer at 3.1Å and 3.5 Å respectively. The hydrogen bond made with LYS 511 became slightly shorter at 2.5Å. TYR 523 made two new hydrogen bond interactions with a hydroxyl group and nitrogen at 2.7Å and 2.5Å respectively. GLU 384 made a new 3.1Å hydrogen bond with Candidate 2. Additionally, a 3.2Å T-shape pi-pi stacking interaction was made between PHE 527 and Candidate 2. The overall more negative docking score was due to the new hydrogen bonds and pi-pi stacking interactions. These new interactions compensate for the weaker bonds and provide a higher binding affinity for the active site.

Similar to the ADMET properties of Moexipril and Candidate 1, Candidate 2 had no known toxicophores or aggregators, and did not violate Lipinski Rule of 5. The molecular weight of the drug decreased to 440.45 g/mol, less than both the original drug and Candidate 1, increasing the chance of being absorbed. The XLogP increased to 1.34 which is lower than Candidate 1 but higher than Moexipril. However, it still falls in the range of oral bioavailability. There were four additional Lipinski hydrogen bond donors compared to Moexipril, increasing the number of possible hydrogen bonds and electrostatic interactions with the active site, creating a higher binding affinity. There are two chiral centers, which is the same as Candidate 1, but less than Moexipril. This decreases the chance of having undesired inactive or toxic effects. Based on the docking score, molecular interactions, and ADMET properties, Candidate 2 is expected to be a better lead drug than Moexipril and Candidate 1.

### Homology Analysis

To test the efficacy and viability of new drug candidates for treating hypertension, preclinical animal studies need to be done to test the safety and toxicity of the drugs. Looking at organisms with homologous proteins is an important step in identifying a good model system for preclinical studies. Using BLAST, we discovered two animals with homologous proteins to the human ACE. *Pan troglodytes* (chimpanzee) and *Mus musculus* (house mouse) have 100% query cover and a high percentage identity. *Pan troglodytes* and *Mus musculus* have percentage similarity identities with human ACE of 99.6% and 81.32% respectively. *Mus musculus* is a good starting organism to test new drug candidates because there is an abundance and are easier to maintain compared to *Pan troglodytes*. We then used Jalview to do an in-depth protein sequence alignment analysis of each homologous enzyme to see if the drug candidates would interact with the active site. The sequence alignment showed that GLN 281, LYS 511, TYR 520, and TYR 523 were conserved in both species which means they will likely interact with Moexipril, Candidate 1, and Candidate 2. PHE 527 was conserved in both species, which interacts with both Candidate 1 and 2. GLU 384 was also conserved in both species which interacts with Candidate 2. With these results, we can hypothesize that the residues are going to have very similar, if not the same, interactions with the active site. However, strong interactions do not necessarily mean that the drug candidates are safe. Full preclinical animal studies still need to be conducted to test the safety and toxicity of these drugs.

## CONCLUSION

Since hypertension is a common issue that affects millions of people around the world, more treatment options need to be developed to solve this problem. By modifying a known drug molecule using computational and chemical intuition methods, two novel drug candidates were proposed. Comparing the results to the existing ACEI Moexipril, Candidate 1 had a much better docking score due to increased molecular interactions with the active site, as well as desirable ADMET properties. Candidate 2 had even better results compared to Candidate 1 and Moexipril. It had improved ADMET properties, the most negative docking score, and the most interactions with the active site. Initial testing of these drug candidates can be done on animal models with proteins homologous to the human ACE such as *Pan troglodytes* and *Mus musculus*. By using animal models the safety, efficacy, toxicity, and off-target effects of these novel candidates can be further evaluated.

## Author Contributions

Research was designed by all authors; all experiments were carried out by J.D.L. The manuscript was written through contributions of all authors. All authors have given approval to the final version of the manuscript.

## ACKNOWLEDGMENT

Research reported in this publication was supported by UC Davis, the National Science Foundation Award Numbers 1827246, 1805510, 1627539, the National Institute of Environmental Health Sciences of the National Institutes of Health (NIH) under Award Number P42ES004699, UC Davis, NIH Award Number R01 GM 076324-11 and the Rosetta Commons. The content is solely the responsibility of the authors and does not necessarily represent the official views of the National Institutes of Health or National Science Foundation. This study was derived from a course based undergraduate research study conducted in Chemistry 130B at UC Davis.

